# WikiGenomes: an open Web application for community consumption and curation of gene annotation data in Wikidata

**DOI:** 10.1101/102046

**Authors:** Tim E. Putman, Sebastien Lelong, Sebastian Burgstaller-Muehlbacher, Andra Waagmeester, Colin Diesh, Nathan Dunn, Monica Munoz-Torres, Gregory S. Stupp, Andrew I. Su, Benjamin M. Good

## Abstract

With the advancement of genome sequencing technologies, new genomes are being sequenced daily. While these sequences are deposited in publicly available data warehouses, their functional and genomic annotations (beyond genes which are predicted automatically) mostly reside in the text of primary publications. Professional curators are hard at work extracting those annotations from the literature for the most studied organisms and depositing them in structured databases. However, the resources don’t exist to fund the comprehensive curation of the thousands of newly sequenced organisms in this manner. Here, we describe WikiGenomes (wikigenomes.org), a web application that facilitates the consumption and curation of genomic data by the entire scientific community. WikiGenomes is based on Wikidata, an openly editable knowledge graph with the goal of aggregating published knowledge into a free and open database. WikiGenomes empowers the individual genomic researcher to contribute their expertise to the curation effort and integrates the knowledge into Wikidata, enabling it to be accessed by anyone without restriction.

## Introduction

Sequencing an organism’s genome has become a routine procedure in the life sciences, but until that genome is annotated, it tells us very little about the organism. The knowledge captured in genomic and functional annotations provides the ‘blueprint’ of the biology of the organism that can be leveraged to drive all manner of scientific inquiry. The knowledge represented by annotations is mostly published in the free text of scientific journal articles. In order to make this type of knowledge computable and accessible as structured annotations, we typically rely on centralized curation efforts like those supported by model organism databases such as the Zebrafish Model Organism database (ZFIN) (1) and Mouse Genome Informatics (MGI) (2). Unfortunately, the vast majority of sequenced genomes do not have a dedicated curation team, hence new curation models need to be explored.

Wikidata is a recently developed project of the Wikimedia Foundation that enables the collaborative construction of a centralized graph database (3). It was initially created as a central hub for structured data across all the language-specific Wikipedias, but its potential applications are much broader. Wikidata follows the Resource Description Framework (RDF) (https://www.w3.org/RDF/ (4)) model, the W3C standard for data interchange. RDF links related entities together into ‘triples’, defined by a subject concept, an object concept, and a predicate that describes the relationship between them. This semantic structure allows the modeling of complex systems in a queryable knowledge graph (3,5). The contents of Wikidata currently represent 28 million concepts, from all domains of knowledge, spanning over 1.3 billion triples (grafana.wikimedia.org/dashboard/db/wikidata-query-service).

While other resources have also mapped their data onto an RDF data model (6–8), Wikidata is unique in that it allows both read and write access. Wikidata content can be both queried and edited programmatically via an application programming interface (API) (www.**wikidata**.org/w/**api**.php) and can be queried using the Wikidata Query Service (WDQS) (query.wikidata.org) via the SPARQL Protocol and RDF Query Language (4,9,10). SPARQL is a query language for graph databases similar in syntax to SQL for relational databases.

Notably, all Wikidata content is in the public domain, eliminating the potential for problematic restrictions on downstream reuse and redistribution (11). It is populated through various programs (“bots”) (12) en masse, and through the discrete contributions of individual users through Wikidata’s own web interface or through a handful of tools developed by Wikimedia Foundation’s WMF Labs (tools.wmflabs.org/).

There is growing appreciation within the bioinformatics community for the potential of Wikidata as an open resource that can be populated, queried and edited by anyone. However, Wikidata has not yet widely reached the biological domain experts who have much to gain and contribute. Taking advantage of the querying capabilities of Wikidata can be a challenge for biologists as structured query languages like SPARQL are not commonly part of a researcher’s toolkit. Further, the domain-specific data models (e.g. the patterns of connections between genes, proteins, and their annotations (www.wikidata.org/wiki/Wikidata:WikiProject_Molecular_biology)) underlying the content in the Wikidata graph are not readily apparent nor automatically enforced by the generic Wikidata web interface. This makes providing meaningful contributions to the graph a challenge for the researcher and a roadblock for community curation in general. While Wikidata offers a powerful technical platform for constructing an open community knowledge base of unprecedented scope, it cannot reach this potential without the large-scale participation of experts in the domains of knowledge that it seeks to represent. To attract specialists, e.g. life scientists, its content must be presented in compelling applications that are more useful than the applications that these specialists currently use to do their work.

Here, we describe WikiGenomes (www.wikigenomes.org), the first domain-specific application built on Wikidata, tailored to the needs of biomedical researchers. WikiGenomes is a Web application that is designed and built to allow the genomic researcher to contribute to curating the knowledge they are discovering. It facilitates both the consumption and community curation of genomic data in Wikidata, extending the reach of the biocuration effort deeper into the long tail of sequenced genomes.

## Results

### Wikidata

Our group and others have been working to populate Wikidata with a comprehensive and centralized knowledge graph that represents biomedical knowledge in a structured, queryable, and computable format (5,12). Previously, we loaded genomic data from a variety of organisms including *Homo sapiens* “human” (www.wikidata.org/wiki/Q15978631), *Mus musculus* “mouse” (www.wikidata.org/wiki/Q83310), *Rattus norvegicus* “brown rat” (www.wikidata.org/wiki/Q184224), *Saccharomyces cerevisiae S288c* “baker’s yeast” (www.wikidata.org/wiki/Q27510868), and *Macaca nemestrina* “Southern pig-tailed macaque” (www.wikidata.org/wiki/Q618026). In this work, we systematically expanded this list to include all genes (390,719) and proteins (372,178) from the 120 NCBI prokaryotic reference genomes (https://www.ncbi.nlm.nih.gov/genome/browse/reference/) (**Figure 1**). These organisms are all reasonably well-studied, but in general did not have community-maintained genome databases. In addition to organism/genetic data, we contribute to and maintain data for proteins, chemical compounds and diseases (5,12,13).

To ensure that the work we are doing in Wikidata is trusted as high quality by the scientific community, our group references the source and provenance of all claims in a consistent and structured way. Direct links to the data sources allow the user to see for themselves the source of an annotation and read further in the referenced publication to come to their own conclusions. S<u>tandards for representing the evidence and pr</u>o<u>venance</u> of any claim made using our infrastructure is outlined in a detailed project wiki (https://www.wikidata.org/wiki/User:ProteinBoxBot/evidence). References can be accessed either manually by the web interface or programmatically via the Wikidata API (https://www.wikidata.org/w/api.php).

In addition to recording data provenance, Wikidata claims can be qualified by how the initial claim was made. For example a Gene Ontology (GO) annotation can be qualified using the Gene Ontology Evidence Codes to indicate the determination method for this claim. The combination of references and qualifiers provides the necessary context to the stated claim, making it more trustworthy and useful. This model allows the researcher to, for example, treat electronically curated annotations differently from those that were manually curated.

### Data Aggregation and Maintenance in Wikidata

As a foundation, we loaded key information for each genome about each gene and gene product into Wikidata. Wikidata is not simply an open database into which data can be dumped. It is a structured knowledge graph that requires careful consideration and community consensus in its design. Currently, these entity models in Wikidata are not comprehensive mirrors of those models in their sources (e.g. NCBI, UniProt, etc…). We have chosen to aggregate key data from a variety of sources, creating a minimal viable model that can be expanded on by us or others. The basic data in Wikidata for all the organisms we have loaded includes genomic position and orientation, Gene Ontology annotations, Enzyme Commission Numbers, InterPro domains and external database identifiers. The data model that we have created in Wikidata, and the sources of the annotations and entities are illustrated and reported in Figure 1.

**Figure 1.**
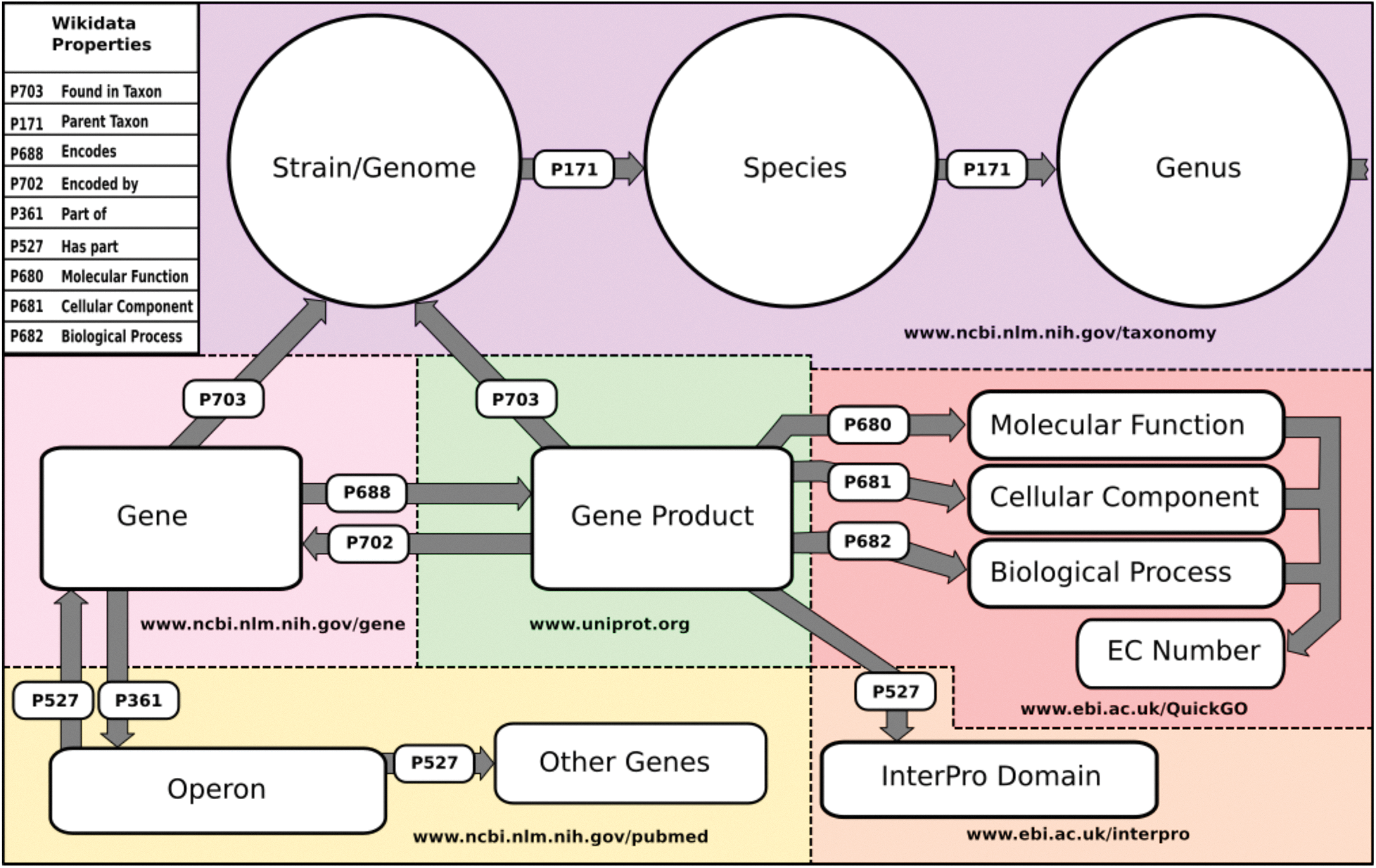
Wikidata Data Model and Sources. Schematic of the basic structure of the data model in Wikidata. Entities include purple: Organism data sourced from NCBI’s taxonomy database (www.ncbi.nim.nih/taxonomy), light pink: gene data sourced from NCBI’s Gene Database (www.ncbi.nim.nih/gene), green: Gene product data sourced from UniProt (www.uniprot.org), red: Gene Ontology annotations sourced from EBI’s QuickGO API (www.ebi.ac.uk/QuickGo), orange: InterPro domain annotations sourced from the InterPro project (www.ebi.ac.uk/interpro) and yellow: Operon data currently sourced from primary publications hosted in PubMed (www.ncbi.nim.nih.gov/pubmed). The names of the Wikidata properties that construct the data model are included in the upper left hand corner.

Data retrieval, standardization, loading and maintenance are done routinely (currently monthly) via our publically available WikidataIntegrator Python package (https://github.com/SuLab/WikidataIntegrator and https://pypi.python.org/pypi/wikidataintegrator) as well as M, and we are working towards custom update schedules that would follow each source’s update schedule. WikidataIntegrator interacts with the Wikidata API to create new or edit existing Wikidata items when appropriate. The data points are mapped to the appropriate “properties” in Wikidata (**Supplemental Table 1**). Mapping these properties to a new semantic model in Wikidata is key to aggregation of data from different sources into a single cohesive graph that represents our collective knowledge of each concept and the relationships between those concepts.

An example of other data we have loaded and maintain is the Gene Ontology (http://www.geneontology.org/), which we have loaded to Wikidata to provide semantic framework for gene product annotations. The structure was represented as a hierarchical tree using the Wikidata ‘Subclass of’ (P279) property. We programmatically gathered professionally curated Gene Ontology annotations from the European Bioinformatics Institute’s QuickGo API (https://www.ebi.ac.uk/QuickGO/), and NCBI’s Gene Database FTP resources (ftp.ncbi.nlm.nih.gov/gene/DATA/gene2go.gz) for all of the annotated gene products in Wikidata (5,12,13). Any gene product item in Wikidata that is annotated with a specific GO term will be linked to that GO term’s Wikidata item via the appropriate Wikidata GO property (e.g. *molecular function P680*) (**Figure 1**). Our methods for loading and maintaining other domains, such as disease and chemical data, are described in our previous works (12,13).

### WikiGenomes

To allow non-developer biologists to interface with the genomic data knowledge graph that we developed in Wikidata, we built WikiGenomes. WikiGenomes currently supports the 120 NCBI Bacterial reference genomes (https://www.ncbi.nlm.nih.gov/genome/browse/reference/) and thier genes and gene products. We designed WikiGenomes to allow a user to consume genomic data from any organism in Wikidata in a single web application. The user begins by selecting an organism using a type-ahead search feature on the application’s landing page. Once an organism has been selected, the application is directed to the primary interface page where the data for the selected organism is loaded and displayed. In addition to providing access to genetic data for these organisms, WikiGenomes also allows users to directly contribute genomic and protein annotations to the Wikidata knowledge graph (**Figure 2**).

**Figure 2.**
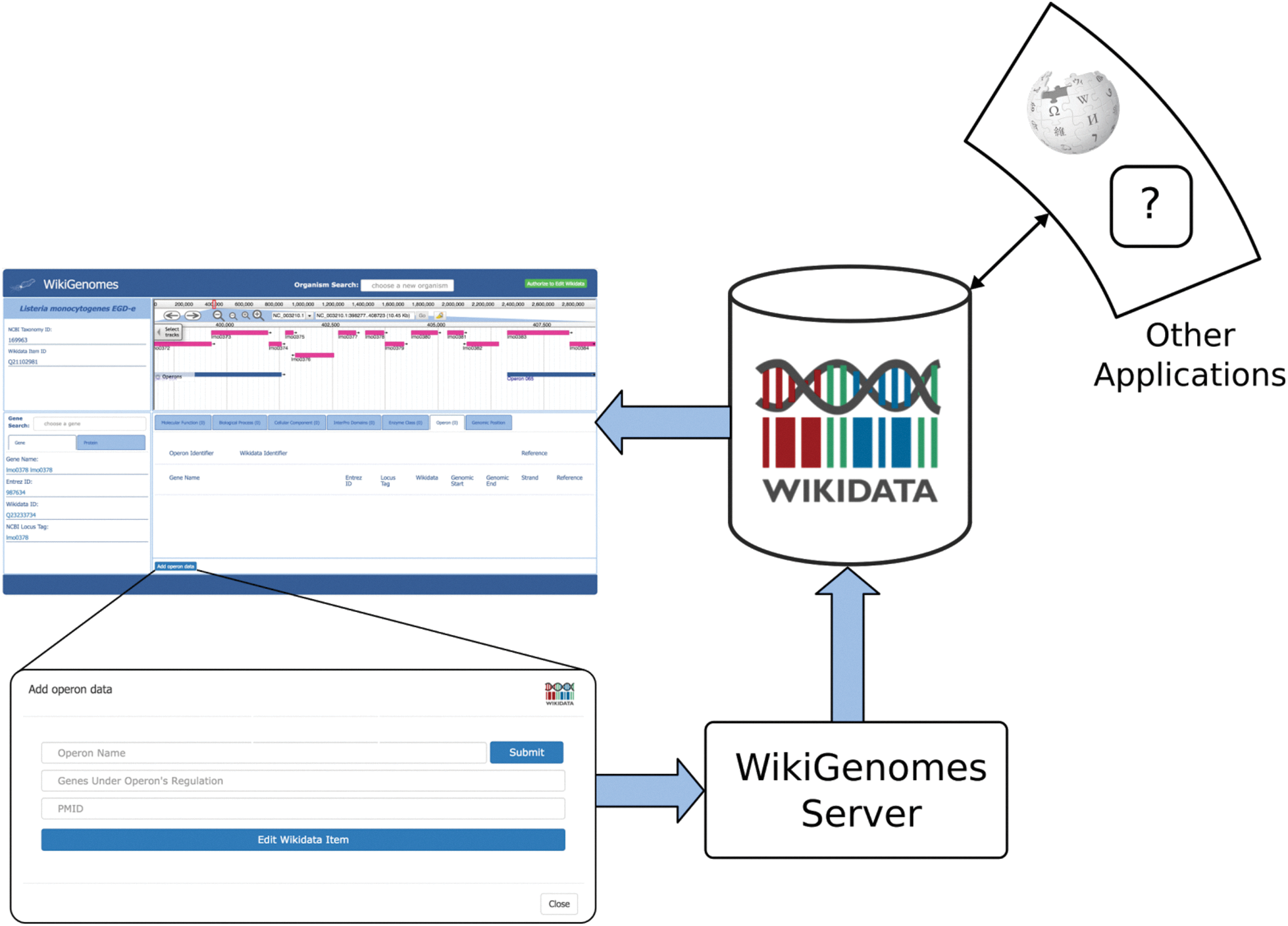
Process of contributing to and consuming data in WikGenomes via Wikidata. WikiGenomes retrieves data from Wikidata via the WDQS SPARQL query service, and allows contribution of defined edits back into Wikidata via a guided annotation curation process. The data then becomes available to any web application that uses Wikidata in a similar way.

The primary page of the interface is divided into four windows that render the data for the selected organism: its genome, genes/proteins, and annotations for the currently displayed gene/protein. To select and load data, there are two forms that allow organism and gene/protein selection (**Figure 3**). Each of the data elements are drawn dynamically from the Wikidata knowledge graph using the WDQS SPARQL query system. The genome data is rendered in the open source JBrowse genome browser (14,15). The WebApollo and JBrowse development teams expanded on already existing code in the JBrowse framework to allow JBrowse to load annotations gathered from Wikidata SPARQL queries to the WDQS endpoint (16). The DNA sequences themselves are curated by the RefSeq Project (16) and because bulk primary data is not suitable for storing in Wikidata, are collected monthly from GenBank’s FTP web service (See **Supplementary File 1** for all Wikidata queries that drive the system and **Supplementary Table 2** for the FTP paths of data collected from NCBI). The open source code for WikiGenomes is available at https://github.com/putmantime/CMOD.Django.

**Figure 3.**
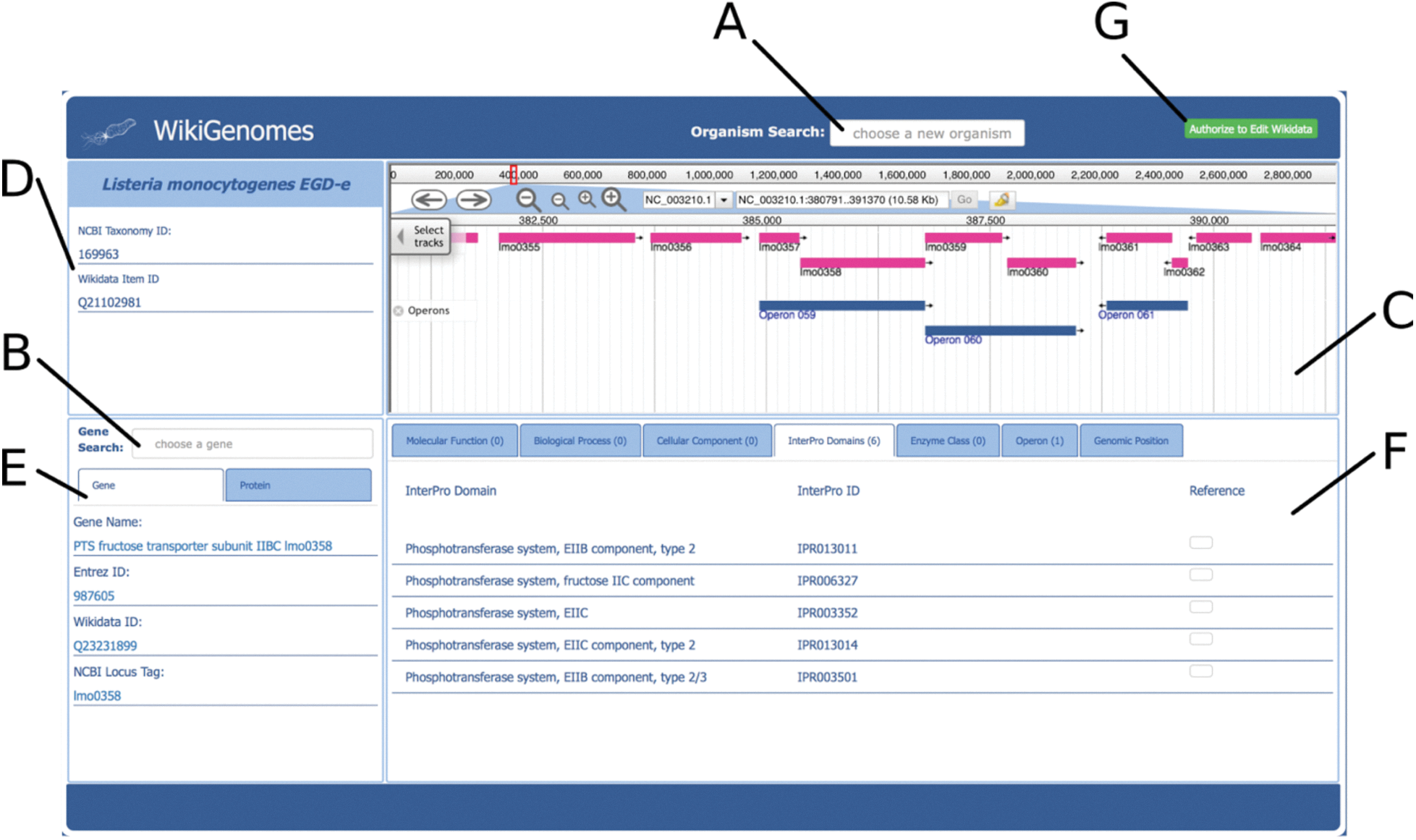
Overview of the WikiGenomes Interface. **A.** The ‘Organism Search’ form selects the organism and populates the main page of the application with data specific to that organism. **B.** The ‘Gene Search’ form loads data for a selected gene/protein. **C. Genome** The WikiGenomes genome browser is an instance of JBrowse, a high-performance, web-based, client-side genome browser that currently displays gene and operon tracks. **D. Organism** The “Organism Box’ displays the name of the selected organism and basic core identifiers (hyperlinked to their respective database entries). These include the Wikidata ‘QID’, and the taxonomy ID from NCBI’s Taxonomy Database. This content window is where any type of metadata about the organism will be added as it becomes available in Wikidata (e.g., morphology, lifestyle, Gram staining, disease associations, drugs that have action against it). **E. Gene/Protein** The ‘Gene/Protein’ content window displays the gene name and basic identifiers for the currently loaded gene and gene product in a tabbed view. The ‘Gene’ tab includes the NCBI Entrez ID, the Wikidata QID, and the NCBI Locus Tag (a core identifier for bacterial genes). The ‘Protein’ tab contains the UniProt ID, NCBI RefSeq Protein ID and Wikidata QID. These are the core identifiers required for the current data model in Wikidata, but as more and more mapped identifiers from different resources are added to Wikidata, they will be displayed here. **F. Annotations** The annotations that have been curated in Wikidata for the currently loaded Gene/Protein are displayed in the content window in the bottom right corner of the application, separated and organized by navigation tabs. The tabs represent the domains of annotations that are pulled from various resources into Wikidata for bacterial genes and proteins. Currently, annotation types include Molecular Function, Biological Process, Cellular Component, InterPro Domain, Operon, Enzyme Commission Number and Genomic Position. Each annotation type displays its own relevant information consisting of hyperlinked database identifiers, the hyperlinked Wikidata identifier, the annotation itself, and (if appropriate) genomic coordinates of the annotation (which are also rendered in the genome browser). All annotations are linked to references retrieved from their underlying Wikidata statements. **G. Authorize to Edit Button.** This button redirects to WikiMedia.org login where the user can user login to their Wikidata account and authorize WikiGenomes to make edits using their credentials.

While bioinformaticians can programmatically gather data and upload it to Wikidata, WikiGenomes provides a domain specific editing interface for non-programmers. WikiGenomes distills editing tasks into straightforward forms that guide the user through the process of building annotations. Internally, the server translates the data entered through these forms into the structures needed to populate the Wikidata knowledge graph and submits them via the Wikidata API (**Figure 2**). WikiGenomes uses the WikiMedia OAuth extension to allow users to edit Wikidata using their own Wikidata account (www.mediawiki.org/wiki/Extension:OAuth). This makes it possible to utilize the Wikidata edit history to track the contributions of individual editors, potentially offering ways to reward good editors as well as mechanisms for detecting vandalism automatically (Good 2012).

### Community Curation Forms

There are currently two editing forms in WikiGenomes, one for adding functional annotations and one for adding operon annotations. The forms are designed to allow a user to curate data from a publication by prompting the user to provide the necessary information and references to make a useful contribution to Wikidata (**Figure 4**).

**Figure 4.**
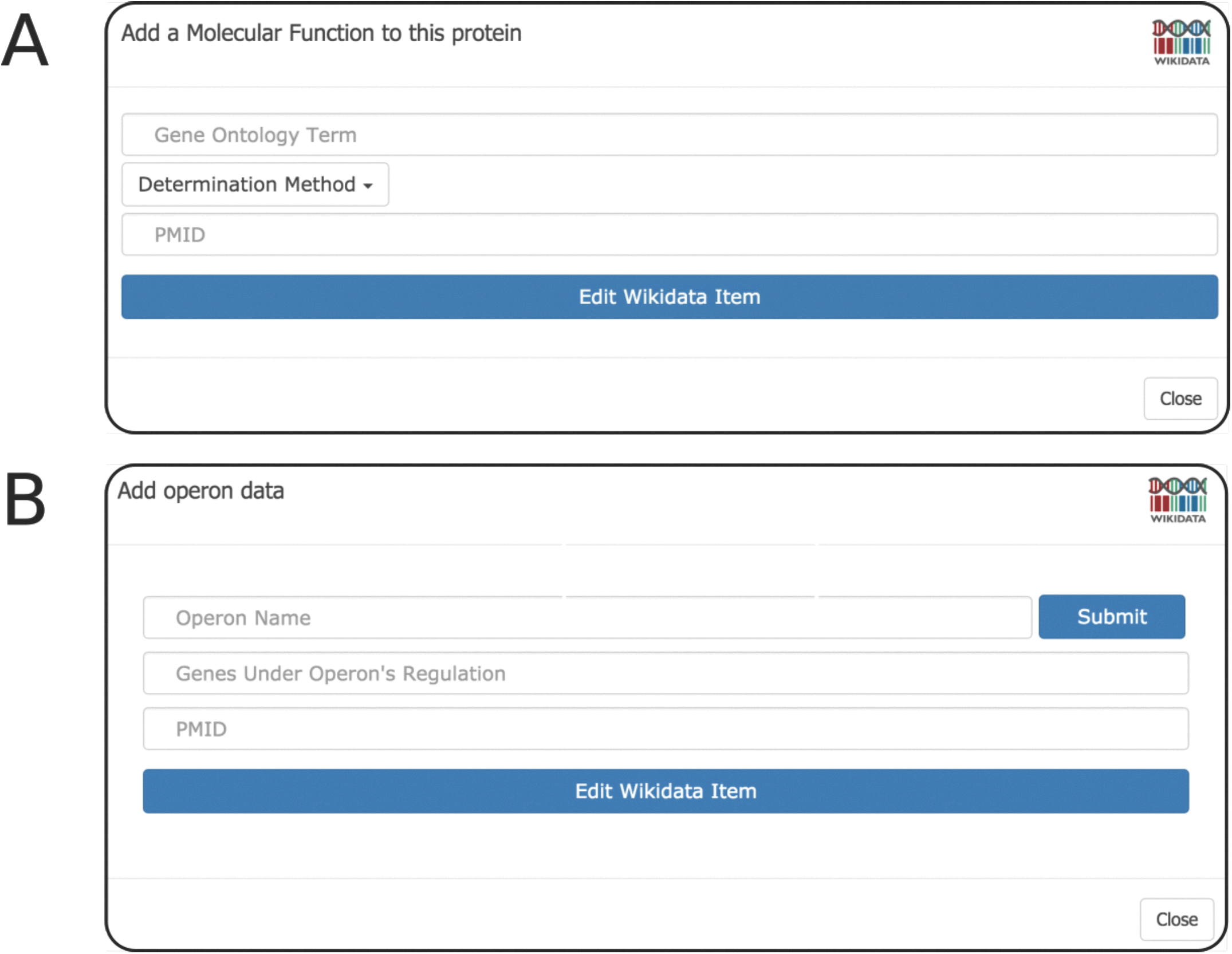
Editing Forms. **A. Gene Ontology Form** The Gene Ontology Form prompts the user to supply 3 pieces of information: 1) the Wikidata item of the GOterm, 2) the method that was used for determination (GO evidence code *i.e. derived from experiment, sequence similarity, etc…*) and 3) the PMID of the publication that the statement originated from. The GO term selection box incorporates type-ahead autocomplete allowing the user to find and select the proper GO term to describe the annotation. Upon user submission, WikiGenomes submits the API call to write the new annotation to Wikidata. **B**. **Operon Form** The Operon Form prompts the user to enter the name of the operon; if the operon already exists, WikiGenomes will find it in Wikidata and add to it. If the operon does not exist in Wikidata, the new name will be set as the label of a new Wikidata item for that operon. The user then provides the genes whose expression is regulated by the operon. The input field for genes also provides type-ahead functionality, allowing the user to quickly search for and select genes from that genome. Like all WikiGenomes forms, the user must also provide the publication that the statement was derived from.

### Gene Ontology Form

Molecular function, cellular component and biological process annotations types are added to Wikidata (and subsequently WikiGenomes) through the Gene Ontology Annotation Form (**Figure 4A**). The WikiGenomes GO form allows users to make structured GO annotations to gene products by identifying an annotation from a primary publication, mapping the annotation to the proper GO term, and then pushing it to Wikidata referencing the publication. We have chosen to adhere to the GO annotation model, rather than user generated free-text, to guide the annotation process in a consistent manner. This is accomplished by using the form to search Wikidata by typeahead for the proper GO term (as mentioned we maintain the GO ontology in Wikidata), selecting the proper GO evidence code (http://geneontology.org/page/guide-go-evidence-codes) to provide a determination method (i.e. EXP: Inferred from Experiment http://geneontology.org/page/exp-inferred-experiment), and select the publication as a reference using the PubMed Identifier (PMID) (**Figure 4A**). When the annotation is submitted, the annotation data is pushed to Wikidata using the Wikidata API and the annotation will be viewable in WikiGenomes shortly after (there is ~5 min delay as the Wikidata SPARQL endpoint updates).

### Operon Form

Regulatory annotations are rare in large data warehouses like NCBI and regulatory annotation curation would greatly benefit from community involvement. To this end we created a form to allow user curation of prokaryotic operons (**Figure 4B**). The WikiGenomes Operon Form works similarly to the GO form in using typeahead search boxes to retrieve relevant data from Wikidata, as well as input boxes to upload novel data. The user can create a new operon or add genes to an existing operon, input the determination method and reference the PMID, then submit that annotation and (as in the GO Form) our backend framework will do the work of creating the structured and standardized annotation in Wikidata.

## Discussion

WikiGenomes was built as a proof a concept that Wikidata can serve as a universal database backend for domain specific applications. WikiGenomes is in the stage of functional prototype and usage stats are thus not yet available. WikiGenomes provides a working demonstration of a new technical pattern that, we posit, offers the potential to fundamentally change how biocuration is organized in ways that, over time, will result in a more efficient movement of knowledge throughout the scientific community. While this is not yet a tool in common usage, we believe that it shows the potential of centralized, community owned knowledge aggregation projects like Wikidata, and hope it will ultimately help stimulate community curation at both the level of individual researchers and bioinformatics laboratories.

### Distributed Curation and Access, Centralized and Integrated Content

The incorporation of operon information for *Chlamydia pneumoniae* into WikiGenomes illustrates the pattern of centralizing content while decentralizing control. Despite prokaryotic operons being heavily studied, the actual annotations are rarely included in genomic assemblies and often reside in supplemental tables of primary publications. *Chlamydia pneumoniae* is a pathogen affecting animals and humans causing lung infections (www.cdc.gov/pneumonia/atypical/cpneumoniae/index.html). The operons of the *C. pneumoniae* strain *CWL029* genome were annotated in a 2011 study (17) but access to this information was only provided in the form of a supplementary Excel file. To our knowledge, none of those ~200 experimentally derived annotations have been curated in GenBank or in operon-specific databases such as DOOR (18) or ODB3 (19). Illustrating the ‘small data to big data’ approach of WikiGenomes, we loaded this content programmatically via the Wikidata API. Individual domain experts could also have loaded these data via the operon form (**Figure 4b**). Using either method, these annotations are now fully-linked data, viewable as a track in the WikiGenomes browser and in any other application drawing content from the open Wikidata knowledge base. Integrating this knowledge into the central Wikidata knowledge graph creates queryable connections to other organisms, diseases, drugs and small molecules. The more extensive the knowledge graph becomes, the more it can be used to explore biological research questions (5),(12,13).

### Stepping Towards Community Curation

So far, the great majority of content accessible through WikiGenomes has been added by our research group. While this content is valuable, it is only a starting point on the path towards an open, community-curated knowledge graph for biology. The fundamental logic of the community curation pattern has been well-described many times (20–22). By incorporating the scientific community into the process of curating the knowledge that they are generating and consuming, we could, in theory, dramatically increase the scale at which biocuration can collectively operate. Here, as usual, the devil is in the details. With certain notable exceptions, most attempts to stimulate community curation fail to attract enough editors to achieve their goals. With the WikiGenomes application, we make no claim that we have solved this challenge of incentives and recruitment. To realize its vision, WikiGenomes will need to transition to a model where the content consumers become the content producers. To make this transition, a number of problems must be addressed. In facing them, some inspiration can be taken from one of WikiGenomes’ most important and successful predecessors, the Gene Wiki effort to organize functional gene information in the context of Wikipedia articles (23).

The key distinction between the Gene Wiki and other projects with very similar goals but less success in recruiting contributors (20,24), was that it was embedded directly in Wikipedia. The Wikipedia context provided the Gene Wiki with immediate discoverability (e.g. Gene Wiki articles rank highly in Google search results), connection to a large community of editors, and a proven social and technical framework for supporting large-scale community content creation. Wikidata now offers most of these same characteristics for structured information. Its content is highly discoverable, with hyperlinks coming in from nearly every Wikipedia article and a high performance query engine that provides a single point of entry into all of its data. It has already attracted thousands of editors from the Wikipedia communities as well as external groups more focused on structured data. It offers a user interface for human editing of content as well as an effective API for programmatic updates. Finally, communities are forming around it in similar ways to Wikipedia. There are social structures in place for administration of the higher-level data structures it utilizes (e.g. the allowed properties) and emerging domain-focused communities such as WikiProject Medicine and WikiProject Molecular and Cellular Biology that work to build and maintain domain-specific content within the larger knowledge graph.

With all of those similarities, one key distinction is that structured information is much more difficult to edit and to present effectively than the hypertext of Wikipedia. While the Wikidata interface (www.wikidata.org) provides a useful, general-purpose foundation for navigating and editing its content, it is not nearly sufficient as a way to connect end-users with the information they need. WikiGenomes provides a genome-focused, editable view of biological knowledge in Wikidata; it offers the large community of scientists an easy way to both access and share knowledge in a structured way, thus opening a door towards open community curation.

### Conclusions and Next Steps

While tools like WikiGenomes are a step in the right direction for converting ‘small data’ to ‘big data’ several key problems remain to be solved in the community curation model. Interfaces like WikiGenomes that provide access to content drawn from openly editable public sources should consider ways to help users assess the trustworthiness of content based on automatic inspection of the provenance information provided in associated references as well as in the histories of the editors who generated the content. In addition to making trust computable, such provenance information will also be of value in attributing the work of the curators who devote time to these open resources - potentially providing an aspect of the solution to the incentive problem. Aside from presenting trust-related information, the forms for adding content should be designed in ways that help domain experts generate high-quality annotations in the first place. The addition of documentation, logical constraints based on underlying ontologies, and helper interfaces are areas where the experience of the existing biocuration resources would be very valuable.

WikiGenomes is a first foray into the world of applications that could be built with the content that is growing in Wikidata. As that content expands and diversifies, many new applications will need to be created to bring that information most effectively to the people that need it and who can contribute to it. Wikidata is unlikely to be the end-all and be-all of open biological knowledge bases. While it has a number of advantages that make it an ideal platform for initiating the endeavor of linking all biological knowledge, it has limitations in terms of content scope that will eventually need to be overcome through the provision of equally open systems that are tailored more specifically to biological content.

Immediate further development of WikiGenomes is focused on teaming up with biocuration groups to help refine the user experience and identify important areas where the data model needs to be expanded. One example would be to work with further with the WebApollo team to integrate WebApollo (25) into WikiGenomes, taking advantage of WebApollo’s annotation features. Another useful feature we will work towards implementing is the ability to export data from Wikidata via WikiGenomes in standard formats (e.g. GFF3). We hope to eventually create a tool in WikiGenomes that brings the biocuration and basic research communities together, creating a more efficient system that will allow for more comprehensive reference genome curation of biological knowledge than is currently possible. Eventually we would make WikiGenomes a deployable toolkit that smaller research communities can customize and deploy while easily sharing their work with the global community.

## Acknowledgements

We would like to thank the Wikimedia foundation for supporting the Wikidata project, and especially the Wikidata community for their efforts in populating it. This project was supported by the National Institutes of General Medical Sciences (R01GM089820) and NIH Common Fund programs for Big Data to Knowledge (U54GM114833) and Extracellular RNA Communication (U54DA036134).

